# CHARM: COVID-19 Health Action Response for Marines – association of antigen-specific interferon-gamma and IL2 responses with asymptomatic and symptomatic infections after a positive qPCR SARS-CoV-2 test

**DOI:** 10.1101/2021.12.13.472352

**Authors:** Martha Sedegah, Chad Porter, Michael R. Hollingdale, Harini Ganeshan, Jun Huang, Carl W. Goforth, Maria Belmonte, Arnel Belmonte, Dawn L. Weir, Rhonda A. Lizewski, Stephen E. Lizewski, Stuart C. Sealfon, Vihasi Jani, Ying Cheng, Sandra Inoue, Rachael Velasco, Eileen Villasante, Peifang Sun, Andrew Letizia

**Affiliations:** Naval Medical Research Center, Malaria Department, 503 Robert Grant Avenue, Silver Spring, Maryland 20910, USA; Naval Medical Research Center, Virology Department, 503 Robert Grant Avenue, Silver Spring, Maryland 20910, USA; Henry M. Jackson Foundation, 6720A Rockledge Dr., Bethesda, MD 20817, USA; GDIT Maryland 20817, USA; U.S. Naval Medical Research Unit SIX, Lima, Perú; Icahn School of Medicine at Mount Sinai, 1 Gustave L. Levy Pl., New York, NY 10029, USA; Leidos, 1750 Presidents Street, Reston, VA 20190, USA

## Abstract

SARS-CoV-2 T cell responses are associated with COVID-19 recovery, and Class I- and Class II-restricted epitopes have been identified in the spike (S), nucleocapsid (N) and membrane (M) proteins and others. This prospective COVID-19 Health Action Response for Marines (CHARM) study enabled assessment of T cell responses in symptomatic and asymptomatic SARS-CoV-2 infected participants.

At enrollment all participants were negative by qPCR; follow-up occurred biweekly and then bimonthly for the next 6 weeks. Study participants who tested positive by qPCR SARS-CoV-2 test were asked to enroll in an immune response sub-study. FluoroSpot interferon-gamma (IFN-γ) and IL2 responses following qPCR-confirmed infection at enrollment (day 0), day 7 and 14 and more than 28 days later were measured using pools of 17mer peptides covering S, N, and M proteins, or CD4+CD8 peptide pools containing predicted epitopes from multiple SARS-CoV-2 antigens.

Among 124 asymptomatic and 105 symptomatic participants, SARS-CoV-2 infection generated IFN-γ responses to the S, N and M proteins that persisted longer in asymptomatic cases. IFN-γ responses were significantly (p=0.001) more frequent to the N pool (51.4%) than the M pool (18.9%) among asymptomatic subjects; however, the difference was not statistically significant (p=0.06) for symptomatic subjects (N pool: 44.4%; M pool: 25.9%). In asymptomatic participants IFN-γ responders to the CD4+CD8 pool responded more frequently to the S pool (55.6%) and N pool (57.1%), than the M pool (7.1%), but symptomatic participants, IFN-γ responses were more frequent to the S pool (75.0%) than N pool (33.3%) and M pool (33.3%). The frequencies of IFN-γ responses to the S and N+M pools peaked 7 days after the positive qPCR test among asymptomatic (S pool: 22.2%; N+M pool: 28.7%) and symptomatic (S pool: 15.3%; N+M pool 21.9%) participants and dropped by >28 days. Magnitudes of post-infection IFN-γ and IL2 responses to the N+M pool were significantly correlated with IFN-γ and IL2 responses to the N and M pools.

These data further support the central role of Th_1_-biased cell mediated immunity IFN-γ and IL2 responses, particularly to the N protein, in controlling COVID-19 symptoms, and justify T cell-based COVID-19 vaccines that include the N and S proteins.

## INTRODUCTION

Coronavirus disease 2019 (COVID-19) is caused by severe acute respiratory syndrome coronavirus 2 (SARS-CoV-2) infection [1], is responsible for more than 220 million confirmed cases, and more than 4.5 million deaths as of September 5, 2021 (COVID-19 report of the World Health Organization). Clinically, SARS-CoV-2 infection ranges from asymptomatic infection to severe illness including acute respiratory distress syndrome (ARDS), and death.

COVID-19 vaccines that induce neutralizing antibodies particularly to the spike (S) protein have been approved for emergency use in the USA, and one, the Pfizer-BioNTech COVID-19 vaccine (Comirnaty) has received U.S. Food and Drug Administration (FDA) approval. However, the efficacy of these vaccines against evolving strains with mutations in the S-protein remains to be fully elucidated [2]. While efforts are ongoing to develop vaccines that incorporate known variant sequences in the next generation S-based COVID-19 vaccines, future vaccines may also need to incorporate other antigenic targets of protective T cell immunity. Other future vaccines involving attenuated whole virus may also be considered. The crucial role of T cell immunity in COVID-19, and persistence of SARS-CoV-2-specific T cell immunity, has critical consequences for vaccine development [3–6]. Antigen-specific T cell responses could support the inclusion of additional antigenic vaccine targets in next generation COVID-19 vaccines. Such vaccines may enhance, broaden, and prolong protective cellular and humoral immunity against COVID-19 by targeting immunodominant regions of multiple antigenic targets and provide T cell-mediated immunity against variants that may escape antibody mediated immunity targeting just the Spike protein.

Since T cells play a vital role in protective immunity against SARS-CoV-2 [7], robust induction of T cell responses by vaccines will be crucial. It is important to understand the range of T cell responses to T cell epitopes of SARS-CoV-2, and *in silico* methods have been used to predict epitopes, often using machine learning [8]. These methods either exploit genetic similarities between SARS-CoV-2 and SARS-CoV particularly for structural proteins S, N, M and E-proteins [9, 10] or peptide-HLA binding prediction methods [8] using artificial neural networks including NetMHCpan derived methods [11–14]. However, many of the predicted epitopes have proven to be non-immunogenic and the accuracy of these predictions can be validated by experimental data [15]. Internal viral proteins are usually more conserved than surface proteins and are often the targets of CD8+ T cells, emphasizing the relative importance of the N-protein among others [16, 17]. Defining a comprehensive set of epitopes enables the breadth of responses and number of epitopes among infected individuals and may help explain heterologous clinical outcomes [7, 18–20].

This study was designed to comprehensively measure IFN-γ cell responses in the COVID-19 Health Action Response for Marines (CHARM) study, a prospective, longitudinal cohort study of healthy, young adults [21]. This cohort enabled the evaluation of immune responses in symptomatic and asymptomatic SARS-CoV-2 infections, as well as in uninfected participants.

## METHODS

### Ethics

The CHARM study was conducted at the Marine Corps Recruitment Depot, Parris Island, South Carolina and has been previously described [21]. Institutional Review Board approval was obtained from the Naval Medical Research Center (protocol number NMRC.2020.0006) in compliance with all applicable US federal regulations governing the protection of human participants. All participants provided written informed consent.

### Study Participants

As previously reported [21], after completing a 14-day home quarantine, recruits who were 18 years or older were eligible to participate. Only those who had a negative qPCR for SARS-CoV-2, at enrollment were considered for this analysis. A 14-question clinical assessment was administered [22], mid-turbinate nares swab specimens obtained for qPCR to detect SARS-CoV-2, and peripheral blood mononuclear cells (PBMCs) were collected at enrollment [22]. Study participants were followed-up on days 7 and 14, 28, 42, and 56 after enrollment at which time they reported symptoms since the last encounter and had qPCR testing for SARS-CoV-2 repeated.

### Study Design

The complete design of this study is shown in Figure 1. Peripheral PBMCs were obtained at enrollment (T0). Additional qPCR testing was performed at days 3/4, 7, 10/11, 14, 28, 42 and 56. Participants with a positive qPCR test were asked to participate in an immune response sub-study. If willing, consented subjects underwent further qPCR tests performed biweekly within the first 14 days after infection, and then bimonthly at 28, 42 and 56 days. Blood was drawn at each of these time points. Samples were categorized into 3 groups: samples obtained in the first week after the initial qPCR positivity (T7), samples obtained in the second week after qPCR positivity (T14), and >15 days after qPCR positivity (long-term, TLT). All participants completed a questionnaire reporting 14 specific COVID-19 related symptoms and were characterized as asymptomatic (124 participants) for those with no symptoms, and symptomatic (105 participants) for those with any reported COVID-19 symptoms.

**Figure 1.**
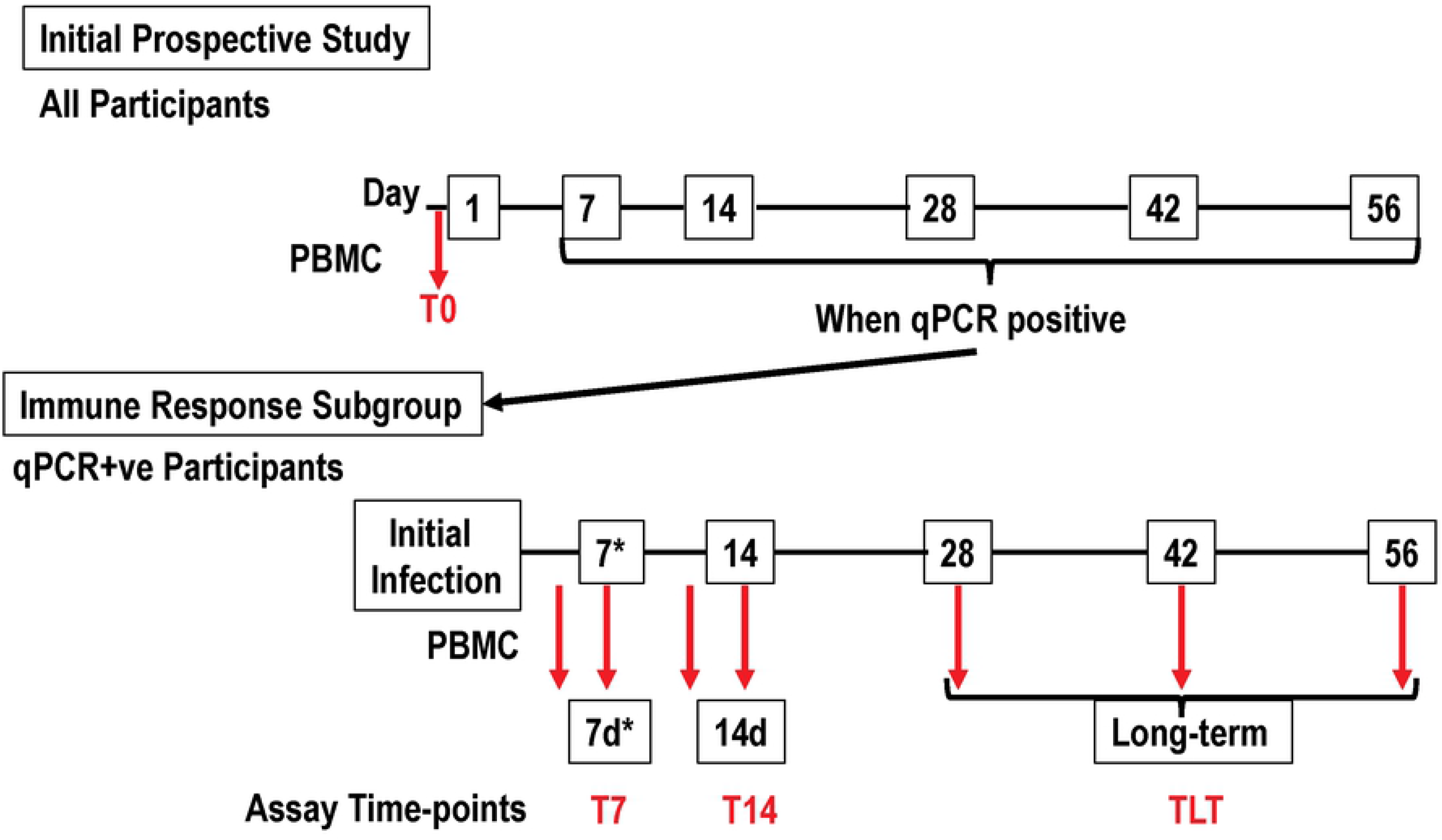
Participant Flow: times of FluoroSpot assays relative to the first positive PCR test. Participants were enrolled in the initial prospective study and tested by qPCR biweekly initially and then bimonthly. When qPCR positive, the participant was transferred to the immune response subgroup within 48-96 hours (7d*) and tested by qPCR biweekly and bimonthly thereafter. PBMCs were isolated (red arrows) prior to the initial qPCR (at enrollment, T0), and once positive at ¾ days 7 days (grouped into T7), 10/11 days (grouped into T14), and greater than 28 days (time long-term (grouped into TLT) post qPCR positivity for SARS-CoV-2.

### Immune samples

PBMCs for measuring cell-mediated immunity (FluoroSpot assay) were collected at the time of enrollment (T0) and then after the participant’s first positive qPCR test (T7, T14, TLT), isolated from heparin tubes within 24 hours, and stored in liquid nitrogen until used. Cryopreserved PBMCs were thawed, washed, counted, and used in the FluoroSpot assay to measure cells secreting either interferon-gamma (IFN-γ), Interleukin-2 (IL2), as previously reported [23–25].

### Peptide pools

All peptides were obtained from BEI Resources (Manassas, VA). The full-length spike glycoprotein (S), full length nucleocapsid (N) protein, and the membrane (M) protein were each covered by a series of 17-mer or 13-mer (aa) peptides overlapping by 10 amino acids that were combined into antigen-specific peptide pools. The S protein pool contained 181 peptides, the N protein pool contained 59 peptides, and the M protein pool contained 31 peptides. In addition, all N and M peptides were combined into a single N+M pool. Samples were tested against the S pool and the N+M pool. Based on availability of cells in some participants, we tested individual antigenic pools of N peptide and M peptide pools. The CD4+ and CD8+ T epitopes peptide pool (kindly provided by Dr. Alessandro Sette, La Jolla Institute for Immunology) [9] and their distribution among SARS-CoV-2 proteins is shown in Supplementary Table S1. CD4+ epitopes were synthesized as 241 15-mers and CD8+ epitopes were synthesized as 628 8-13-mers that were combined into one CD4+ and CD8+ peptide pool [9]. Due to limited PBMCs, not all participants were tested with the CD4+CD8 peptide pool.

### Interferon-gamma/IL2 FluoroSpot assay

Antigen-specific circulating PBMCs were evaluated using pre-coated FluoroSpot plates and kits purchased from Mabtech (Mabtech AB, Nacka Strand, Sweden) and used according to the manufacturer’s instructions. The previously described *ex vivo* ELISpot was modified [26]; briefly, 2-3 x 10^5^ PBMCs suspended in 100 μL complete medium were incubated in the FluoroSpot plates with antigen-specific peptide pool at final concentration of 2 ug/mL suspended in 100 μL complete medium. CTL-CEF-Class I Peptide Pool Plus (Cellular Technology Ltd, Cleveland, OH) consisting of 32 peptides corresponding to defined HLA class I-restricted T cell epitopes from cytomegalovirus, Epstein-Barr virus and influenza virus was used as an internal control for each participant. PHA, a mitogen, was used as a positive control for cell viability. Negative control unstimulated PBMCs received medium only. Cultures were incubated for 40-42 h at 37°C in 5% CO_2_. Each PBMC sample was assayed in duplicate and the number of single-staining antigen-specific IFNγ- and IL2-secreting cells and double-staining IFNγ- and IL2-secreting cells were recognized as spot-forming cells (sfcs) and enumerated using an automated FluoroSpot reader (AID iSpot, Autoimmun Diagnostika GmbH, Strasberg, Germany). After subtraction of the mean number of sfcs in negative control wells (no antigen), the mean of the net sfcs of the test sample was expressed as sfcs/10^6^ PBMCs. A positive response to each individual peptide pool was defined as positive when there was a statistically significant difference (p=<0.05) between the average of the sfc in test and negative control wells (Student’s two tailed *t*-test), plus at least a doubling of sfc in test compared to control wells, plus a difference of at least 10 sfc between text and control wells [27].

### Statistical Analysis

Comparisons of the proportion of participants demonstrating a response were made using a Pearson chi-square while comparisons on the proportion of participants demonstrating responses across pools were made using a McNemar’s Test. Comparisons across study groups were made using a Wilcoxon Rank Sum Tests and comparisons before and after infection were made use paired Wilcoxon Rank Sum Tests. Spearman Rank Correlations were used to assess the correlation between maximum responses post-infection. For each participant, the single highest magnitude of response among the 4 time points was selected for that participant that were defined as greatest number of sfc/106 PBMC at T7, T14 or TLT after the initial SARS-CoV-2 qPCR positive result. All statistical analyses were interpreted using a two-tailed alpha=0.05 and were made using SAS v9.4 or JMP v 12 (SAS Institute; Cary, NC). In some cases, asymptomatic and symptomatic participants who all had responses to the N+M or CD4+CD8 peptide pools were compared for responses to individual S, N or M peptide pools. In the Supplemental Figures, comparisons of the magnitudes of positive responses after the first qPCR assay were analyzed using the Mann-Whitney U-Test.

## RESULTS

### Frequency of IFN-γ and IL2 responses to S, N+M, and CD4+CD8 peptide pools before and after infection in the asymptomatic and symptomatic groups

The individual frequencies of IFN-γ and IL2 responses in asymptomatic and symptomatic participants to the S, N+M, and CD4+CD8 pools at T0, T7, T14 and TLT are shown in Supplementary Figures S1 and S2. The frequencies of IFN-γ and IL2 responses to S and N+M pools significantly (p=<0.001) rose at 7 days after infection and persisted unchanged at TLT (>28 days). The exceptions were IFN-γ response to the S protein that significantly declined by TLT among symptomatic but were unchanged among asymptomatic participants, and IL2 responses to the S pool significantly rose at T7 and again at T14 in the asymptomatic group, but not in the symptomatic group. There were no IL2 responses to the CD4+CD8 pool in both groups, except one symptomatic participant who had a positive response.

We next determined the frequency of positive responses at T0, T7, T14 and TLT to the S, N+M and CD4+CD8 peptide pools.

#### S and N+M peptide pools

among all participants, the frequency of positive IFN-γ responses to the S pool (30.0%) or the N+M pool (37.0%), was higher among asymptomatic (34.4%, 39.2%, respectively) than symptomatic (24.8%, 34.3%, respectively) participants; however, only IL2 responses to S pool were significantly higher (p=0.04) (Table 1A) in the asymptomatic group. IFN-γ responses to the N+M pool were significantly more frequent than to the S pool (p=0.03), whereas there was no difference in IL2 responses (Table 1B).

**Table 1.**
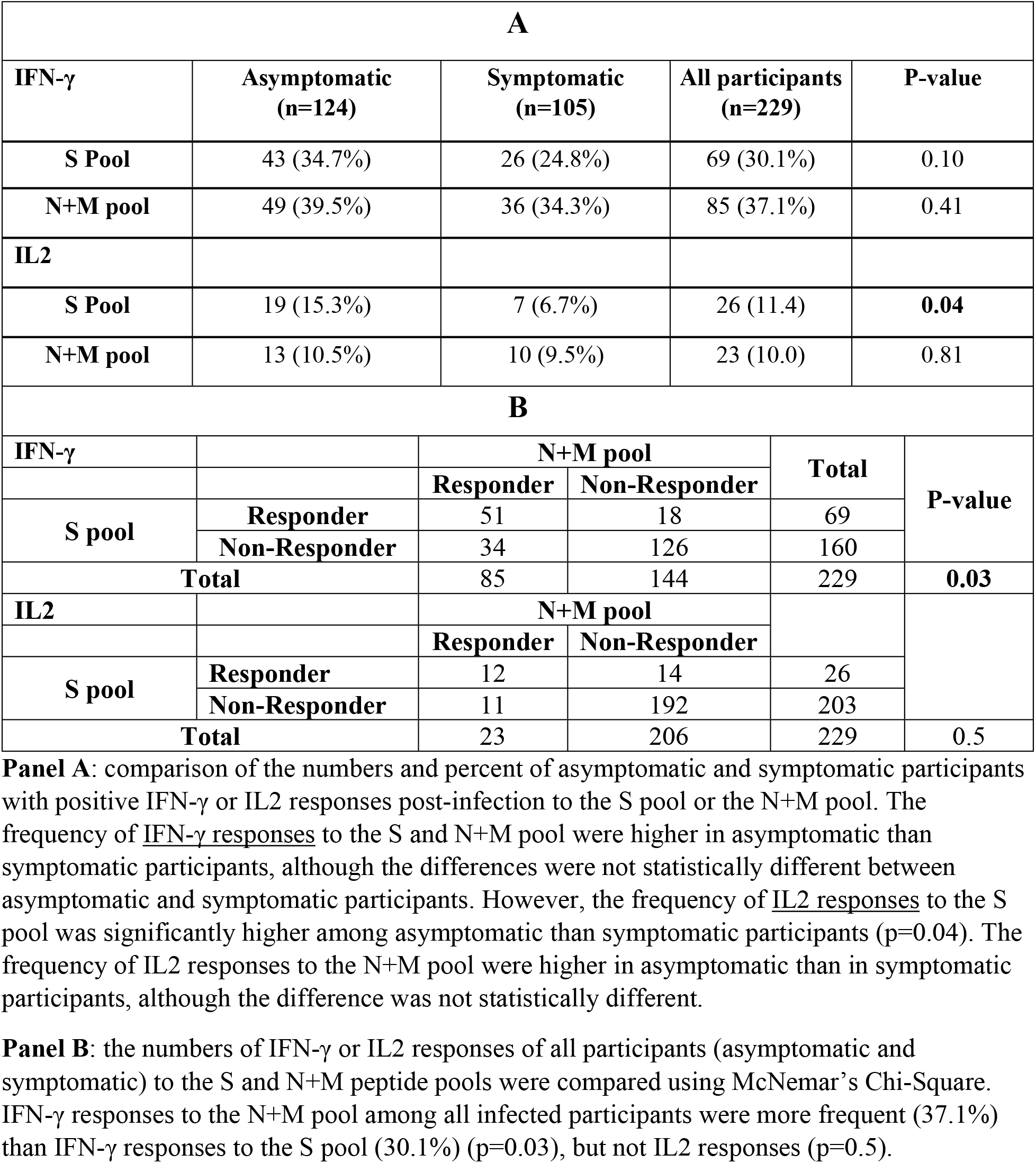
Frequency of immune responses to the S and N+M peptide pools in asymptomatic and symptomatic participants.

We next determined whether the frequencies of IFN-γ and IL2 responders to the N+M pool were associated with the individual N or M pools in asymptomatic and symptomatic participants (Table 2). Among all participants, IFN-γ responses to the N pool (48.4%) were significantly (p=0.002) higher than to the M pool (21.9%). The frequency of IFN-γ responders to the N pool (51.4%) was significantly (p=0.001) higher than the M pool (18.9%) in asymptomatic participants (p=0.001) but was not significantly different among symptomatic participants (p=0.06). The frequencies of IL2 responses were low compared to IFN-γ responses in each group.

**Table 2.** Frequency of immune responses to the N+M, N, and M peptide pools in post-infection asymptomatic and symptomatic participants.

| **Cytokine** | **Pool** | **Asymptomatic**<br>**N+M pool responders** | **Symptomatic**<br>**N+M pool responders** | **All participants**<br>**N+M pool responders** |
| --- | --- | --- | --- | --- |
| **IFN-γ** | **N pool** | 19/37 (51.4%) | 12/27 (44.4%) | 31/64 (48.4%) |
|  | **M pool** | 7/37 (18.9%) | 7/27 (25.9%) | 14/64 (21.9%) |
|  | **p-value** | **0.001** | 0.06 | **0.002** |
| **IL-2** | **N pool** | 2/9 (22.2%) | 0/8 (0%) | 2/17 (11.8%) |
|  | **M pool** | 2/9 (22.2%) | 1/8 (12.5%) | 3/17 (17.7%) |
|  | **p-value** | 1.0 | -- | 0.6 |

The numbers and percent positive of asymptomatic and symptomatic participants with positive IFN-γ or IL2 responses post-infection to the N+M pool were compared with responses to the individual N or M pools.

#### CD4+CD8 peptide pool

among CD4+CD8 pool responders, the frequency of positive IFN-γ in matched asymptomatic and symptomatic post-infection to S, N, and M pools were compared. The responses to the S pool (63.3%) were higher than the N (47.8%) or M (17.4%) pools. However, in asymptomatic participants IFN-γ responses to S and N pools were similar (55.6% and 57.1%, respectively) and higher than to the M pool (7.1%) (Table 3), whereas in symptomatic participants, IFN-γ responses to the S pool were higher (75.0%) than to the N and M pools (33.3% each) (Table 3). This suggests that among asymptomatic participants in this study with IFN-γ responses to the CD4+CD8 pool, responses are generally directed to epitopes within the S pool and N pool and not within the M pool, whereas in symptomatic participants, IFN-γ to the CD4+CD8+ responses are more frequent to epitopes within the S protein than the N protein.

**Table 3.** Frequency IFN-γ and IL2 responses to CD4+CD8 pools and S, N, and M pools in matched asymptomatic and symptomatic post-infection participants.

| **Cytokine** | **Pool** | **Asymptomatic**<br>**CD4+CD8 pool**<br>**responders** | **Symptomatic**<br>**CD4+CD8 pool**<br>**responders** | **All participants**<br>**CD4+CD8 pool**<br>**responders** |
| --- | --- | --- | --- | --- |
| **IFN-γ** | **S pool** | 10/18 (55.6%) | 9/12 (75.0%) | 19/30 (63.3%) |
|  | **N pool** | 8/14 (57.1%) | 3/9 (33.3%) | 11/23 (47.8%) |
|  | **M pool** | 1/14 (7.1%) | 3/9 (33.3%) | 4/23 (17.4%) |
| **IL-2** | **S pool** | Low | Low | Low |
|  | **N pool** | Low | Low | Low |
|  | **M pool** | Low | Low | Low |

The numbers and percent positive of asymptomatic and symptomatic participants with positive IFN-γ or IL2 responses post-infection to the CD4+CD8 pool were compared in matched participants with responses to the individual S, N, or M pools. Low responders had activities that did not meet the positivity criteria (Methods).

### Magnitude of IFN-γ and IL2 responses to S, N+M and CD4+CD8 peptide pools before and after infection in the asymptomatic and symptomatic groups

We next compared the magnitudes of the maximum post-infection responses to each peptide pool in asymptomatic and symptomatic participants. As with determining frequencies of responses, we used the largest magnitude of responses at T0, T7, T14 or TLT.

#### S and N+M peptide pools

The magnitude of IFN-γ and IL2 responses to the S pool were significantly higher in asymptomatic than symptomatic participants (p= 0.03 and p=0.002, respectively); however, only IL2 responses to the N+M pool were significantly higher in asymptomatic participants (p=0.04) (Figure 2). Yet, the magnitude of IFN-γ responses to the N+M pool among all infected participants was higher than the magnitude of responses with S pool (Table 4, p=0.01).

**Figure 2.**
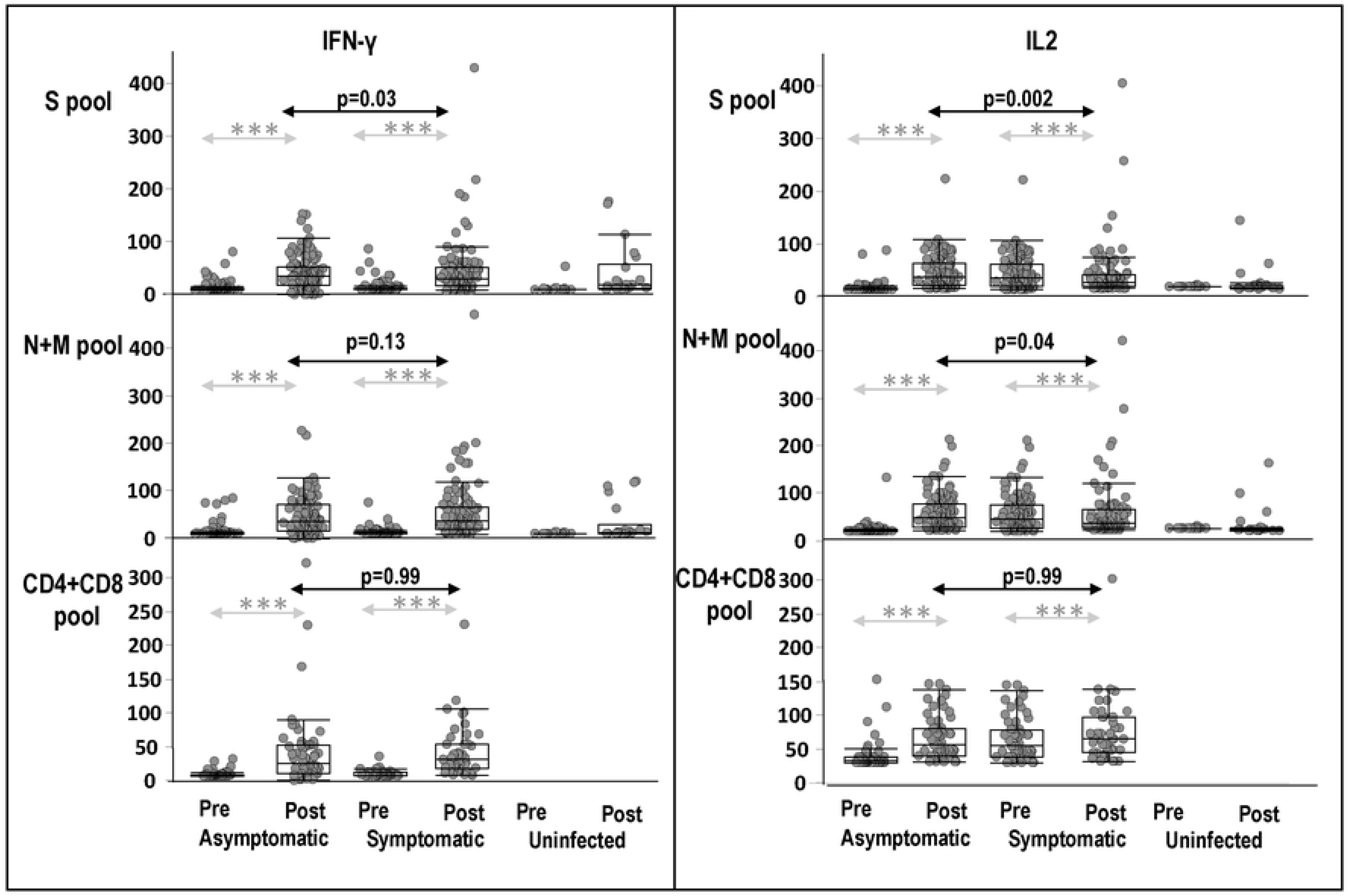
Magnitudes of IFN-γ and IL2 responses of asymptomatic, symptomatic participants before and after infection, and healthy uninfected participants, to S, N+M, and CD4+CD8 pools. Pre-infection (Pre) and maximum IFN-γ and IL2 responses post-infection (Post) to S, N+M, and CD4+CD8 peptide pools of asymptomatic and symptomatic participants, and healthy uninfected participants (Pre = baseline and Post = post-baseline). Significance of differences between Pre and Post *** p=<0.001.

**Table 4.** Magnitude of IFN-γ and IL2 responses to the S and N+M peptide pools among all infected participants (N=229)

| **IFN-γ** | **Mean (Std. Dev.)** | **Median (Q1, Q3)** | **P value\*** |
| --- | --- | --- | --- |
| **S pool** | 40.4 (43.4) | 30.0 (13.3, 52.5) | **0.01** |
| **N+M pool** | 46.7 (50.7) | 32.5 (15.0, 67.5) |  |
| **IL2** |  |  |  |
| **S pool** | 19.3 (26.5) | 12.5 (3.8, 27.5) | 0.2 |
| **N+M pool** | 17.7 (23.4) | 10.0 (3.8, 22.5) |  |

Responses were expressed as box plots: Std. Dev.: standard deviation; Q1: Quartile 1; Q3: quartile 3; \*Signed Rank Test.

Significant correlation (using Spearman Rank Correlations) was observed between maximum post-infection IFN-γ and IL2 responses to the N+M pool and the N pool and the M pool (Figure 3). Maximum IFN-γ responses were more strongly correlated to the N pool (Spearman R=0.67, p<0.0001) than to the M pool Spearman R=0.49, (p<0.0001) suggesting the relative importance of responses to the N pool in the responses to the N+M pool.

**Figure 3.**
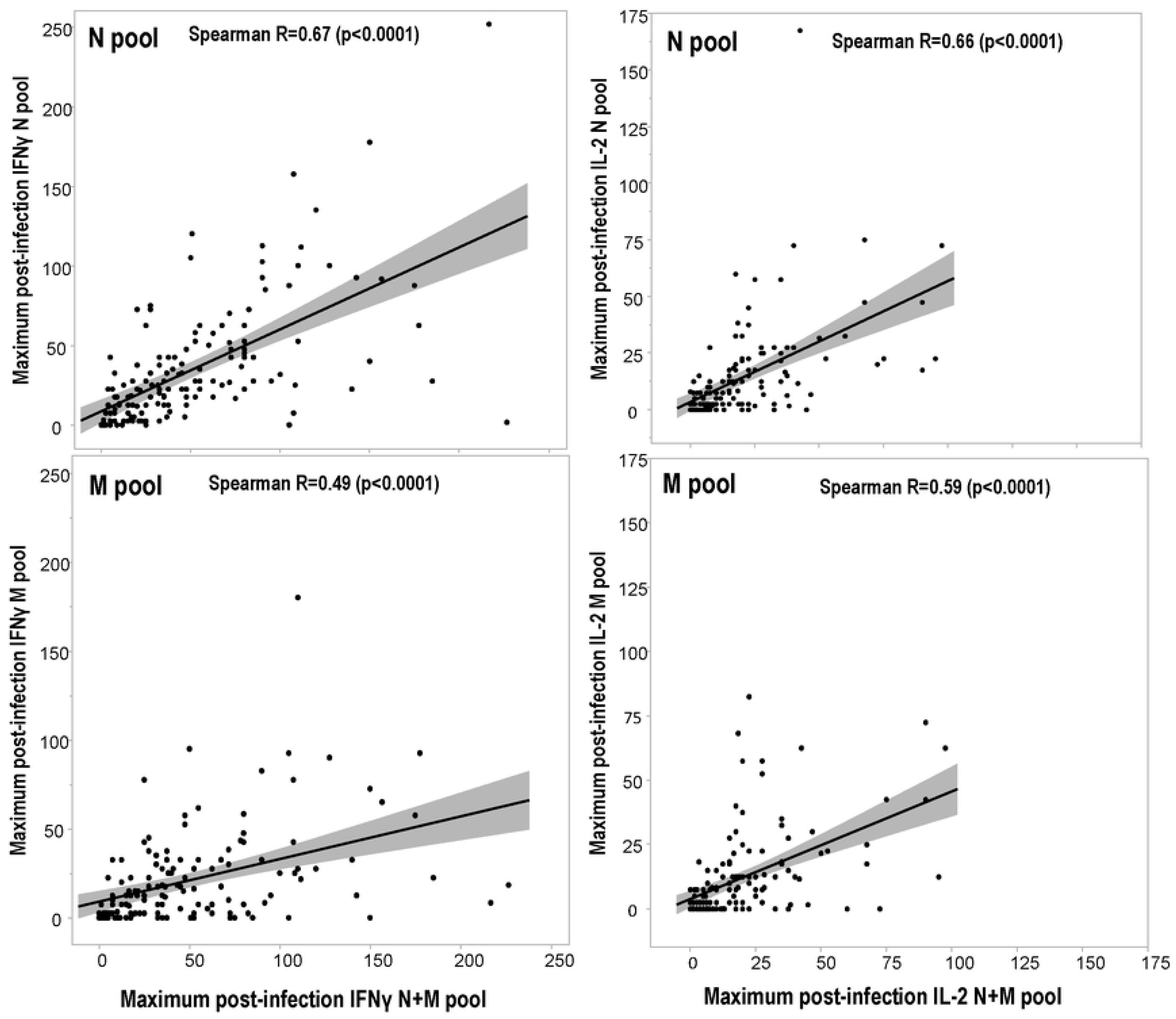
Correlations of the magnitude of IFN-γ and IL2 responses post-infection to the N and M pools with IFN-γ and IL2 responses to the N+M pool. The maximum IFN-γ and IL2 responses post-infection to the N and M pools were compared to those to the mixture of N+M pool in matched participants by Spearman Rank Correlations.

#### CD4+CD8 peptide pool

there as a significant correlation (using Spearman Rank Correlations) between the maximum post-infection IFN-γ responses to the CD4+CD8 pool and post-infection IFN-γ responses to the S, N and N+M pools (comparisons excluded the M pool alone group) (Figure 4). These correlations suggest that that responses to the S and N proteins contribute to the CD4+CD8 epitope pool responses, while the contribution of responses to the M pool alone remain undetermined, and we cannot exclude responses to other non-S, N and M proteins in the CD4+CD8 pool. In asymptomatic participants, CD4+CD8 pool responses are generally directed to epitopes within the S protein and N protein and not within the M protein, whereas in symptomatic participants, CD4+CD8+ responses are more frequent to epitopes within the S protein than the N protein (Table 3).

**Figure 4.**
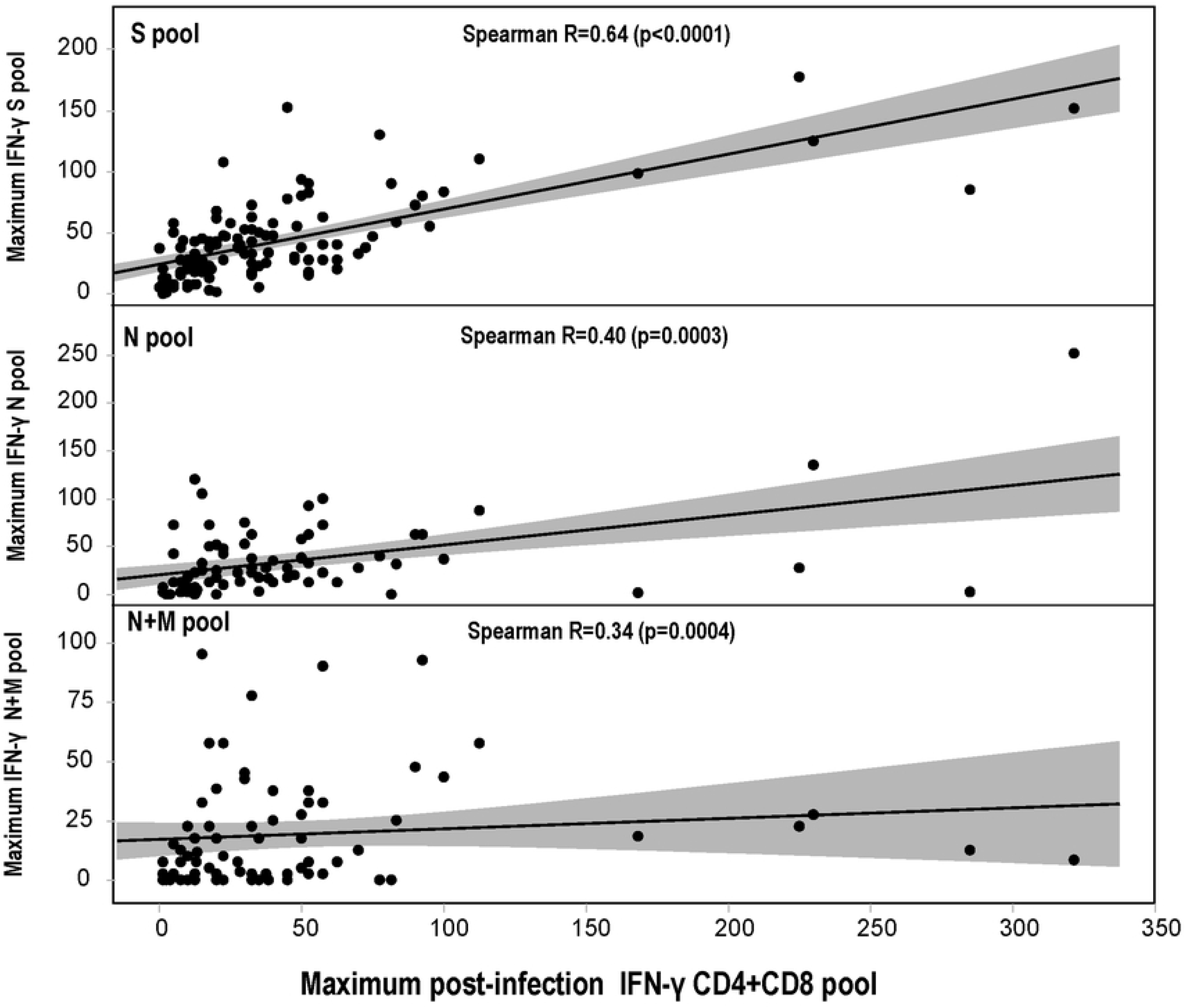
Correlations of the magnitude of IFN-γ responses post-infection to the S, N, and N+M pools with IFN-γ responses to the CD4+CD8 pool. The magnitudes of IFN-γ responses to the CD4+CD8 pool were compared to the magnitudes of IFN-γ responses to the S, N, and N+M pools. Using Spearman’s ranked correlations, IFN-γ responses to the S, N and N+M pools were significantly correlated.

## DISCUSSION

This study enabled comparisons of asymptomatic and symptomatic immune responses immediately after a positive SARS-CoV-2 qPCR assay. We used recovered cells from cryopreserved samples that may not reflect total peripheral T cell counts. Both groups developed frequent and robust IFN-γ responses, although IL2 responses were lower. Others have previously suggested that a significant virus-specific T cell response was not associated with disease severity [7, 28-30], and that total peripheral T cell counts were reduced in asymptomatic patients with reductions in CD4+ and CD8+ T cell counts [31, 32] or greater reductions in CD8 cell counts [31, 33]. Our results suggest that the specificity of responses, notably to the N protein, may be an important indicator of disease status, as symptomatic participants had lower responses to the N protein. We found that asymptomatic participants developed IFN-γ and IL2 responses primarily directed toward the N pool and CD4+CD8 pools, and IFN-γ responses to the N pool were more frequent than to the S pool, whereas symptomatic participants developed responses primarily to the S pool, with lower frequencies of responses to the N, M and CD4+CD8 pools. However, we only assessed asymptomatic and symptomatic participants with mild disease who were all treated as outpatients. Therefore, we cannot extrapolate our findings to other studies using patients with severe disease that suggested lung-homing T cells may contribute to immunopathology, while non-suppressive SARS-CoV-2-specific T cell responses may limit pathogenesis and promote recovery from severe COVID-19 [29].

Early induction of IFN-γ SARS-CoV-2-specific T cell responses to the S protein has been previously associated with mild disease and accelerated viral clearance [28, 30] and the magnitude of responses were more robust in patients with mild disease, whereas those responses were less pronounced in one patient with fatal disease [30]. Using peptide pools in the ELISpot IFN-γ assay, 95% of donors with mild disease had T cell responses to at least one antigen [34]; median aggregate responses were higher in donors with symptomatic disease than asymptomatic disease, confirming another report using ELISpot IFN-γ [4], and was correlated with peak antibody responses. When CD4+ and CD8+ responses were measured, IL2 responses were dominant in the CD4+ subset. However, there was a greater proportion of CD8+ T cells than CD4+ T cells in mild cases [4]. In convalescent patients, using predicted HLA class I-restricted epitopes, there were distinct patterns of immunodominance for CD4+ and CD8+ T cells, accounting for over 80% of the response, confirming an earlier more limited study [9], and CD4+ and CD8+ responses were highly correlated. Our study extends these findings to include the role of the N protein and CD4+CD8+ T cell responses.

While there are no clearly-defined immune correlates of protection against COVID-19 [35], there is considerable evidence that neutralizing antibodies, an elevated CD8+ T cell response and TH1-biased CD4+ effector responses provide optimal protective immunity [36]. Recent results from convalescent COVID-19 participants indicate that CD4+ and CD8+ T cell responses were similar across different SARS-CoV-2 variants [37]. Thus, it is important to better understand the roles of antibody and T cell responses, including IFN-γ and IL2, after vaccination [38], including in the hybrid immunity of subjects that were infected and then subsequently immunized with a COVID-19 vaccine [39–42], as well as their roles in multisystem inflammatory syndrome in children (MIS-C) and post-acute sequelae SARS-CoV-2 infection (PASC) [43, 44].

A major concern of current antibody-based vaccines are mutations in the S protein that affect sequences recognized by vaccine-induced neutralizing antibodies [45–51]. However, it has been suggested that it is highly unlikely that SARS-CoV-2 mutations would affect T cell immunity as so many SARS-CoV-2 epitopes are distributed throughout virus [52, 53]. However, mutations in new variants, including the Delta variant, significantly reduced T cell responses to epitope peptides in convalescent and vaccinated subjects [51], and in *in vitro* binding assays [54]. The genetic HLA-restriction of T cell responses associated with disease outcomes would greatly advance our understanding of the relative importance of predicted epitopes including those used in this study, and whether newly emerging variants of concern carry mutations within these protective epitopes. For example, the L452R mutation contributes to evasion of HLA-A24-mediated cellular immunity [55].

Finally, we observed T cell IFN-γ, but not IL2, responses to the S and N+M pool in some uninfected participants, suggesting cross-reactive T cells derived from prior exposure to the common cold human coronaviruses that is in agreement with previous studies that have reported SARS-CoV-2 cross-reactive CD4+ T cells in unexposed people [56, 57], and others have speculated whether these may contribute to disease outcomes in COVID-19 [58].

Taken together, our results support previous findings on the potential importance of the N-protein for vaccine development [59]. Immunization with the N protein protects against SARS-CoV-2 in non-human primates [60] and mice [61]. Another advantage to considering the vaccine potential of the N protein is its sequence conservation between SARS-CoV-2 and SARS-CoV and MERS-CoV [62]. The N protein contains a region of T cell cross-reactivity that is common to human alpha and betacoronaviruses, as well as a dominant B cell epitope [63].

### Limitations

There are several limitations to this study. Firstly, the study population is young, healthy adults with few comorbidities and may not be reflective of the general population limiting the generalizability of the findings. Secondly, all illnesses were mild, limiting the ability to assess a range of clinical outcomes. Thirdly, the timing of FluoroSpot responses may not have been optimal and further post-infection analyses are warranted. Fourthly, we used cryopreserved cells, and it is possible the recovery of viable cells after thawing may have varied and affected assay readouts. Finally, although we tested immune responses to the S, N and M proteins, it is likely that responses to other proteins may also have a significant role in disease modulation, and responses may partially reflect the relevant sizes of each protein and the numbers of peptides in each pool that were higher for the S pool. We only measured IFN-γ and IL2 producing cells and it is also possible that different cytokines would help to better understand disease severity [30], and phenotypic analysis would also better identify the roles of CD4+ and CD8+ T cell responses.

## ACKNOWLEDGEMENTS

The authors thank Alessandro Sette, Daniela Weiskopf and Alba Grifoni of the Division of Vaccine Discovery, La Jolla Institute for Immunology, La Jolla, CA, who contributed greatly to the success of this project. We also thank Corey Balinsky and Qi Qiu, Naval Medical Research Center, Silver Spring, MD, for their valued technical assistance. MS, CP, EV, and PS are employees of the U.S. Government; CG, DLW, RAL, SEL and AGL are military service members. This work was prepared as part of their official duties. Title 17, U.S.C., §105 provides that copyright protection under this title is not available for any work of the U.S. Government. Title 17, U.S.C., §101 defines a U.S. Government work as a work prepared by a military Service member or employee of the U.S. Government as part of that person’s official duties. The views expressed in this article are those of the authors and do not necessarily reflect the official policy or position of the Department of the Navy, the Department of Defense, nor the U.S. Government.

---

**Who Did What:**

| NAME | Contribution |
| --- | --- |
| Martha Sedegah | Conceptualization, Investigation, Supervision, Formal Analysis, Writing rev/edit |
| Chad Porter | Statistical analysis, formal analysis, visualization, Writing rev/edit |
| Michael R. Hollingdale | Analysis, Writing Original draft, Writing Rev/Edit |
| Harini Ganeshan | Investigation, Analysis |
| Jun Huang | Investigation, Analysis |
| Carl W. Goforth | Investigation, Project Admin, Funding Acquisition, Resources |
| Maria Belmonte | Investigation, Analysis |
| Arnel Belmonte | Investigation, Analysis |
| Dawn L. Weir | Investigation |
| Rhonda A. Lizewski | Supervision, Investigation, Project Administration |
| Stephen E. Lizewski | Supervision, Investigation |
| Stuart C. Sealfon | Funding Acquisition, Project Coordination, Project Administration |
| Vihasi Jani | Investigation |
| Ying Cheng | Investigation |
| Sandra Inoue | Investigation, Analysis |
| Rachel Velasco | Investigation, Analysis |
| Eileen Villasante | Resources, Writing Rev/Edit |
| Peifang Sun | Conceptualization, Formal Analysis, Writing rev/edit |
| Andrew Letizia | Conceptualization, Funding, Supervision, Methodology, Resources, Writing Rev/Edit |

**Table S1:** Distribution of predicted CD4+ and CD8+ epitopes within SARS-CoV-2 proteins.

|  | Number of peptides predicted within protein |  |  |  |  |  | Total |
| --- | --- | --- | --- | --- | --- | --- | --- |
| T cell type | S protein | N protein | M protein | E protein | NS protein | Other Proteins |  |
| Number CD4 Epitopes/protein | 20 | 7 | 8 | 8 | 101 | 97 | 241 |
| % CD4 Epitopes/Total | 8.30 | 2.90 | 3.32 | 3.32 | 41.91 | 40.25 |  |
| Number CD8 Epitopes/protein | 86 | 13 | 15 | 2 | 227 | 285 | 628 |
| % CD8 Epitopes/Total | 13.69 | 2.07 | 2.39 | 0.32 | 36.15 | 45.38 |  |

CD4+ and CD8+ epitopes within each SARS-CoV-2 protein were predicted [52], synthesized, and combined in the CD4+CD8 peptide pool.

**Figure S1: Asymptomatic participants: magnitude and percent positivity of IFN-γ and IL2 responses to the S, N+M and CD4+CD8 peptide pools**

*** p=<0.001; * p=<0.05

The numbers of samples at each time point were based on the numbers of available samples among the 124 asymptomatic participants. Immune responses (sfc/m) and percent positive participants are shown before the first positive qPCR test (T0), 7 days (T7), 14 days (T14) and long-term (TLT) after that first positive PCR test (infection). Immune responses to S, N+M and CD4+CD8 pools were significantly (p=<0.001) higher at T7, and were not significantly different between T7, T14 and TLT after infection; the exception was the significant rise in IL2 responses to S pool between 7d and 14d after infection. There were no positive IL2 responses to the CD4+CD8 pool.

**Figure S2: Symptomatic participants: magnitude and percent positivity of IFN-γ and IL2 responses to the S, N+M and CD4+CD8 peptide pools**

***p=<0.001; **p=<0.01

The numbers of samples at each time point were based on the numbers of available samples among the 105 symptomatic participants. Immune responses (sfc/m) and percent positive participants are shown before the first positive PCR test (T0), and 7 days (T7), 14 days (T14) and long-term (TLT) after that first positive PCR test (infection). Immune responses to S, N+M and CD4+CD8 pools were significantly (p=<0.001) higher at T7, and IFN-γ responses to S protein significantly dropped by TLT. IL2 responses to the CD4+CD8 pool were absent, except in one participant at 14d after infection.

